# Histidine-Triad Hydrolases Provide Resistance to Peptide-Nucleotide Antibiotics

**DOI:** 10.1101/2020.03.04.977132

**Authors:** Eldar Yagmurov, Darya Tsibulskaya, Alexey Livenskyi, Marina Serebryakova, Yury I. Wolf, Sergei Borukhov, Konstantin Severinov, Svetlana Dubiley

## Abstract

The *Escherichia coli* microcin C (McC) and related compounds are potent Trojan-horse peptide-nucleotide antibiotics. The peptide part facilitates transport into sensitive cells. Inside the cell, the peptide part is degraded by non-specific peptidases releasing an aspartamide-adenylate containing a phosphoramide bond. This non-hydrolyzable compound inhibits aspartyl-tRNA synthetase. In addition to the efficient export of McC outside of the producing cells, special mechanisms evolved to avoid self-toxicity caused by the degradation of the peptide part inside the producers. Here, we report that histidine triad (HIT) hydrolases encoded in biosynthetic clusters of some McC homologs or by stand-alone genes confer resistance to McC–like compounds by hydrolyzing the phosphoramide bond in toxic aspartamide-adenosine, rendering them inactive.

**IMPORTANCE:** Uncovering the mechanisms of resistance is a required step for countering the looming antibiotic resistance crisis. In this communication, we show how universally conserved histidine triad hydrolases provide resistance to microcin C – a potent inhibitor of bacterial protein synthesis.

## INTRODUCTION

Microcin C (McC) is a ribosomally synthesized post-translationally modified peptide (RiPP) antibiotic produced by some strains of *Escherichia coli*. Homologous compounds are encoded by gene clusters in numerous gram-negative and gram-positive bacteria (1). McC is produced by *E. coli* cells harboring a conjugative plasmid containing the *mccABCDEF* cluster (2). The *mccA* gene encodes a seven amino acid precursor peptide whose C-terminal residue is modified by the product of the *mccB* gene to yield a peptidyl-adenylate McC^1120^ in which the C-terminal aspartamide is linked to adenosine monophosphate (AMP) through a non-hydrolysable N-acyl phosphoramidate linkage (Fig. 1) (3). McC^1120^ is further modified by MccD and the N-terminal domain of MccE protein, whose joint action results in a fully matured microcin C, McC^1177^, harboring an aminopropyl decoration on the phosphate moiety (Fig. 1) (4). Both forms of McC are exported from the producing cell by a specialized transporter encoded by the *mccC* gene (5).

**Figure 1.**
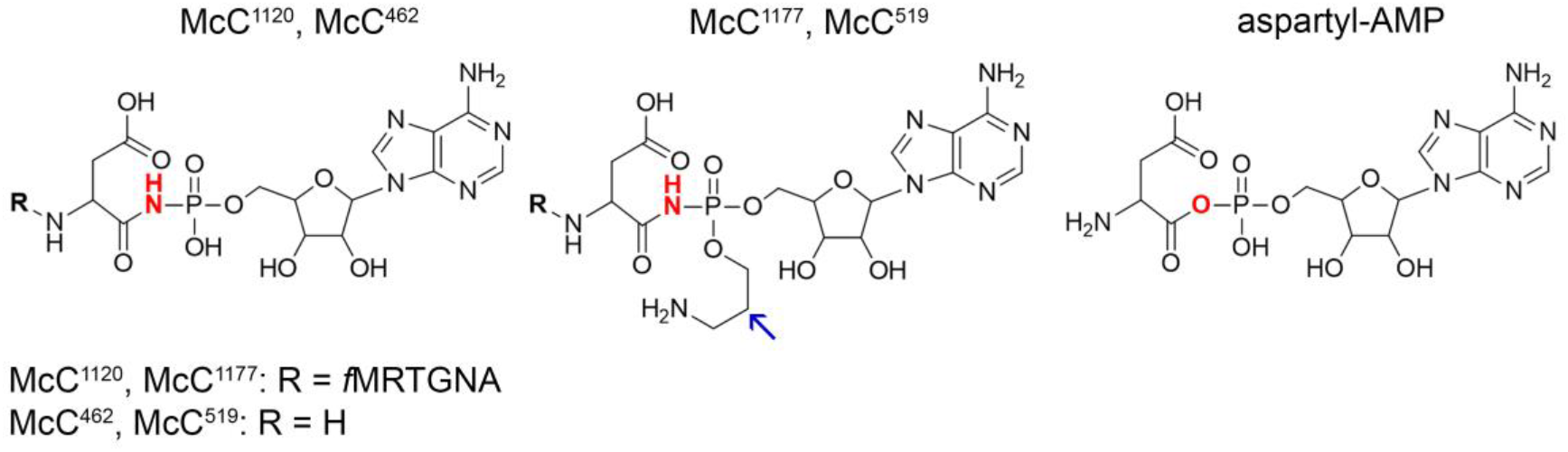
Structures of non-aminopropylated (McC^1120^) and aminopropylated (McC^1177^) forms of *E. coli* microcin C and their processed forms (McC^462^ and McC^519^, respectively), and of Asp-AMP, an intermediate of AspRS-catalyzed reaction. The aminopropyl group is indicated by an arrow.

McC acts through a Trojan-horse mechanism. The peptide part facilitates uptake into the susceptible cell; once inside the cell, the peptide part is proteolytically degraded by aminopeptidases, releasing toxic “processed McC”, a non-hydrolyzable aspartamide-adenylate (Fig. 1), a structural mimic of intermediate of reaction of aminoacylation of tRNA^Asp^ catalyzed by aspartyl-tRNA synthetase (AspRS) (6, 7). Processed McC competitively inhibits AspRS, bringing protein biosynthesis to a halt (5).

Although most McC is efficiently exported outside of the producing cell by the MccC pump, intracellular processing by aminopeptidases should inevitably lead to the accumulation of toxic non-hydrolyzable aspartamide-adenylate and self-intoxication of the producer, since MccC does not export processed McC. Many *mcc*–like clusters acquired additional genes whose products help avoid self-intoxication. In the case of *E. coli*, the C-terminal domain of MccE, a GNAT-type acetyltransferase, acetylates the α-amino group of processed McC making it unable to bind to AspRS (8). In addition, MccF peptidase cleaves the carboxamide bond between the C-terminal aspartamide and AMP of both intact and processed McC (9).

In this work, we report a novel pathway of McC inactivation by Histidine Triad (HIT) superfamily hydrolases encoded in some *mcc*-like biosynthetic clusters or by standalone genes located elsewhere in bacterial genomes. Proteins of HIT superfamily form two separate functional groups: the first group belongs to nucleotide hydrolases, represented by HinT (10–13), Fhit (13), APTX (14), and Dcsp (15) enzymes, while the other group includes nucleotide transferases such as GalT (16). The most ubiquitous members of HIT superfamily – HinT proteins were shown to possess phosphoramidase activity (17). We show that bacterial MccH, a product of a gene in an *mcc-*like cluster from *Hyalangium minutum*, as well as its homologs from *Salmonella enterica*, *Nocardiopsis kunsanensis*, and *Pseudomonas fluorescens*, are phosphoramidases that confer resistance to McC-like compounds by hydrolyzing the toxic aspartamide-adenylate that is produced after intracellular processing of peptidyl-nucleotides.

## RESULTS

### Bioinformatic prediction and experimental validation of an unusual *mcc* operon of *Hyalangium minutum*

Bioinformatics analysis reveals a uniquely organized cluster in the genome of gram-negative bacteria *Hyalangium minutum* DSM 14724 that may determine the production of two putative McC-like compounds. The cluster contains two genes coding for putative precursor peptides, MccA_1_ and MccA_2_, two *mccB* genes, encoding THIF-like adenylyl transferases, and three genes whose products likely constitute a complex ABC-type transporter with integrated HlyD-like translocator and C39-like peptidase (18) (Fig. 2A). An additional gene, *mccH*, is located downstream of the *mccB*_*1*_ gene and encodes a protein belonging to a histidine triad (HIT) superfamily (12).

**Figure 2.**
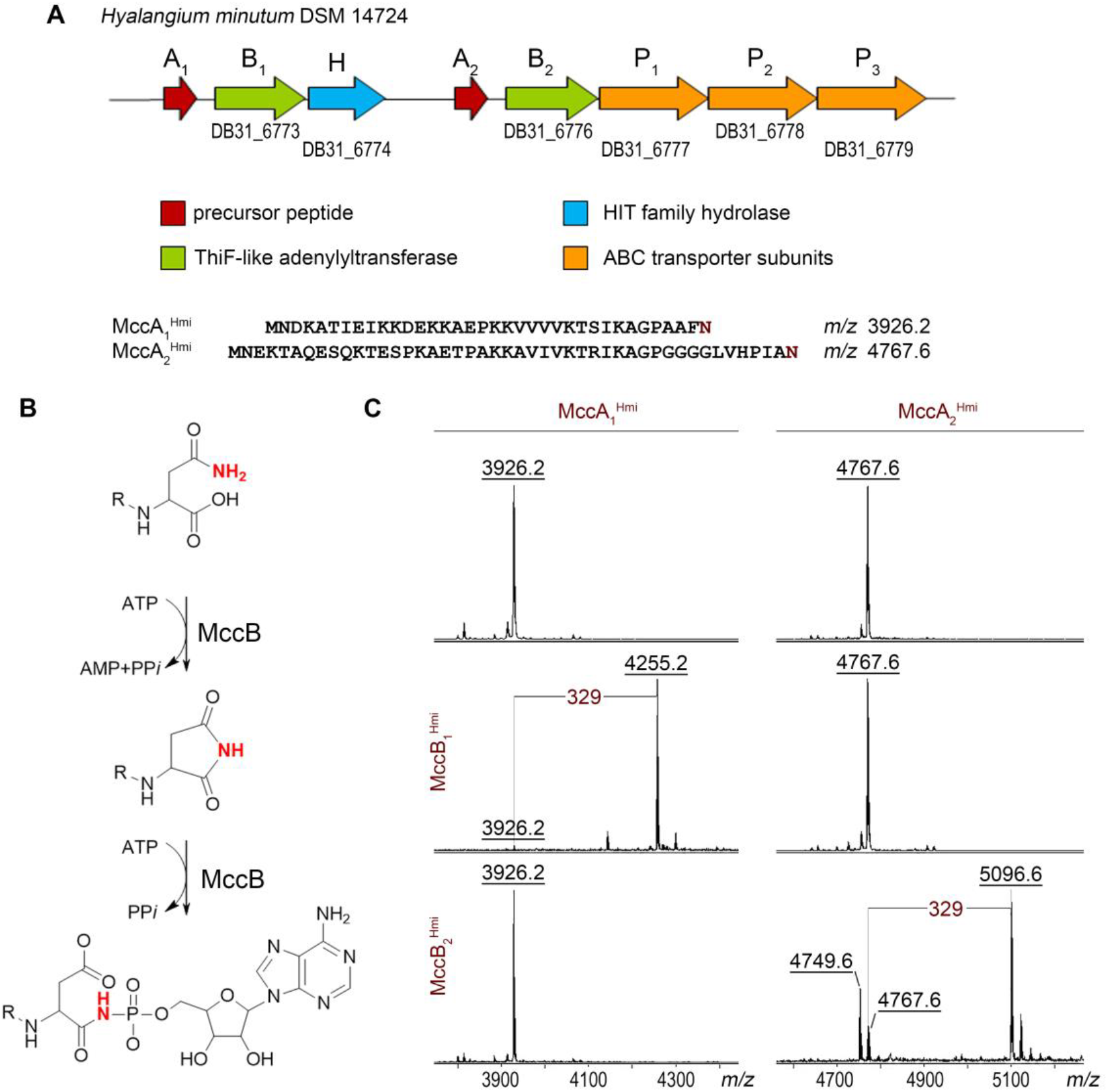
The *mcc*-like operon of *H. minutum* DSM 14724 and its products. (A) Organization of *mcc*-like gene cluster from *H. minutum* DSM 14724. Genes are represented by colored arrows with functional predictions corresponding to each color shown below. (B) The mechanism of MccB-mediated adenylation of an MccA precursor peptide (3). (C) MALDI-TOF-MS spectra of products of *in vitro* reactions between chemically synthesized *H. minutum* MccA_1_ and MccA_2_ precursor peptides and recombinant MccB_1_^Hmi^ and MccB_2_^Hmi^ enzymes in the presence of ATP. Spectra shown at the top are controls (no MccB enzyme added). The mass difference of 329 Da between MccA_1_ (*m/z* 3926.2) and MccA_2_ (*m/z* 4767.6) mass-ions and the mass ions of the reaction products (*m/z* 4255.2 and *m/z* 5096.6, correspondingly) matches modification with AMP; the MH^+^ ion at *m/z* 4749.6 present in reactions containing MccA_2_^Hmi^ and MccB_2_^Hmi^ is a succinimide intermediate of the nucleotidyl transfer reaction.

To validate the predicted *H. minutum mcc*-like cluster, *in vitro* adenylation reactions of synthetic MccA_1_ and MccA_2_ peptides with recombinant MccB_1_ and MccB_2_ adenylyltransferases were performed and products analyzed by MALDI-TOF-MS. As can be seen from Figs. 2B and 2C, incubation of 36 aminoacid-long MccA_1_ (MH^+^ at *m/z* 3926.2) but not 46 aminoacid-long MccA_2_ (MH^+^ at *m/z* 4767.6) with MccB_1_ and an equimolar mixture of four nucleotide triphosphates led to the appearance of a mass-ion at *m/z* 4255.2. The 329 Da mass increase corresponds to the adenylated form of the peptide. Conversely, in the presence of MccB_2_, a prominent MH^+^ ion at *m/z* 5096.6 was observed in reactions containing MccA_2_, corresponding to its adenylated form. An MH^+^ ion at *m/z* 4749.6 with 18 mass units less than the original MccA_2_ ion was also detected. It corresponds to MccB-catalyzed adenylation reaction intermediate – a succinimide derivative of the MccA_2_ peptide. MccA_1_ was not modified by MccB_2_. We conclude that the *H. minutum mcc*-like cluster directs the synthesis of two peptidyl-adenylates, each synthesized from separate precursors by dedicated MccB enzymes.

We next attempted to reconstruct the production of each of the *H. minutum* McC-like compounds in a heterologous *E. coli* host. Cognate *mccA-mccB* pairs were cloned on one expression plasmid, and the *mccP*_*1*_*P*_*2*_*P*_*3*_ genes encoding the putative transporter were cloned on a compatible plasmid. In conditions of induction of plasmid-borne genes, *E. coli* cells or cellular extracts harboring both plasmid pairs did not inhibit the growth of McC-sensitive *E. coli* tester strain and no mass-ions corresponding to *H. minutum* McC-like compounds were detected in cultured medium (Supplementary Fig. S1A). To test if peptidyl-adenylates are synthesized but fail to export from the heterologous host, induced cells were subjected to MALDI-TOF-MS. An MH^+^ ion at *m/z* 4255.2 and 5096.6 corresponding to full-length adenylated MccA_1_ and MccA_2_, respectively, were identified in cells harboring plasmids producing MccA_1_/MccB_1_ and MccA_2_/MccB_2_ pairs. In addition, MH^+^ ions at *m/z* 4124.2 and 4965.6, corresponding to adenylated MccA_1_ and MccA_2_ peptides lacking the first methionine residue, were detected. Unmodified full-sized MccA_1_ and MccA_2_ polypeptides (MH^+^ ions at *m/z* 3926.2 and 4767.6, respectively), and their derivatives lacking methionine (MH^+^ ions at *m/z* 3795.2 and 4636.6) were also observed (Supplementary Fig. S1B). Thus, two distinct products of the *H. minutum mcc* cluster are produced in the heterologous host but fail to be exported at detectable level.

### MccH^Hmi^ confers McC-immunity when overproduced in *E. coli*

We hypothesized that MccH^Hmi^ is a HIT superfamily phosphoramidase that provides *H. minutum* with self-immunity through the inactivation of McC-like compounds. To check this conjecture, we cloned the *mccH*^*Hmi*^ gene in an arabinose-inducible *E. coli* expression vector. We also created plasmids overproducing HinT^Hmi^, a product of a standalone *H. minutum* gene, and its *E. coli* homologue HinT^Eco^. Since the final toxic aspartate-adenylate form of McC produced by *H. minutum* and *E. coli* should be identical (Fig. 1, McC^462^), we tested the susceptibility of MccH- or HinT-expressing cells to *E. coli* McC^1120^. Additionally, we used an aminopropylated “nonhydrolyzable” form of *E. coli* McC, McC^1177^. In the assay, the drops of solutions of two active forms of *E. coli* McC were deposited on lawns of *E. coli* B McC-sensitive cells producing HIT proteins or harboring control empty vector (Fig. 3). The results revealed that the size of growth inhibition zones around drops of fully mature McC^1177^ solution on lawns of HIT protein-producing cells was the same as on control cell lawn. In contrast, *E. coli* cells overexpressing the *mccH*^*Hmi*^ gene were completely resistant to McC^1120^, an intermediate of *E. coli* McC maturation process that does not contain the aminopropyl moiety. The expression of either *hinT*^*Eco*^ or *hinT*^*Hmi*^ had no effect on the size of growth inhibition zones produced by McC^1120^ (Fig. 3). All HIT proteins were produced in comparable amounts, as judged by SDS PAGE (Supplementary Fig. S2). We, therefore, conclude that MccH^Hmi^ but not HinT proteins tested can provide resistance to externally added toxic peptidyl-adenylate.

**Figure 3.**
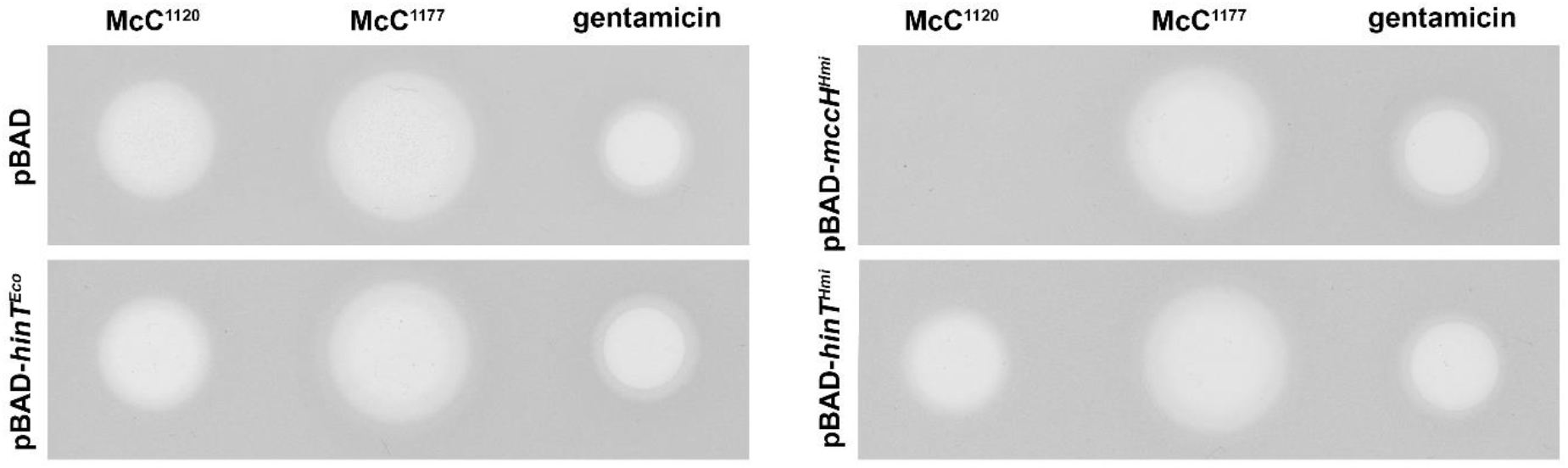
Overproduction of MccH^Hmi^ makes *E. coli* resistant to McC^1120^ but not to mature *E. coli* microcin C McC^1177^. 3 μL of 5 μM solutions of McC^1177^, McC^1120^, or 0.5 μg/mL gentamycin used as control were deposited on lawns of *E. coli* cells harboring indicated plasmids. Results of overnight growth at 37 °C in conditions of induction of plasmid-borne genes are shown.

### MccH^Hmi^ hydrolyzes the phosphoramide bond connecting the aminoacyl and nucleotide moieties of processed McC^1120^

To determine the mechanism of MccH^Hmi^-mediated resistance to toxic peptidyl-nucleotide, the recombinant MccH^Hmi^, HinT^Hmi^, and HinT^Eco^ were purified, incubated with unprocessed McC^1120^ or McC^1177^, and reaction products analyzed by reverse-phase HPLC (RP-HPLC) and MALDI-TOF-MS. No changes were observed after a 1-hour incubation (data not shown). Since no hydrolytic activity against either form of *E. coli* McC was detected, we considered whether the processed forms of McC^1120^ and McC^1177^ could be the substrates of MccH^Hmi^. To this end, aspartamide-adenylates with (McC^519^) and without (McC^462^) the aminopropyl group, were prepared by *in vitro* processing of McC^1177^ and McC^1120^ (see Materials and Methods). After incubation with MccH^Hmi^, HinT^Hmi^, or HinT^Eco^, samples were analyzed by RP-HPLC and MALDI-TOF-MS. Since the expected phosphoramidase activity of MccH^Hmi^ should result in the appearance of AMP or adenosine 5’-phosphoramidate, we used AMP as a marker (Fig. 4A). Upon 1-hour incubation with MccH^Hmi^, McC^462^ was completely converted into a new compound with the same chromatographic mobility as AMP (Fig. 4A). The MH^+^ ion of this compound had *m/z* 348.1, matching AMP (Fig. 4B). The MS-MS fragmentation spectra confirmed this assignment (Fig. 4B). None of the enzymes was able to hydrolyze fully processed microcin with aminopropyl decoration (McC^519^) (Supplementary Fig. S3).

**Figure 4.**
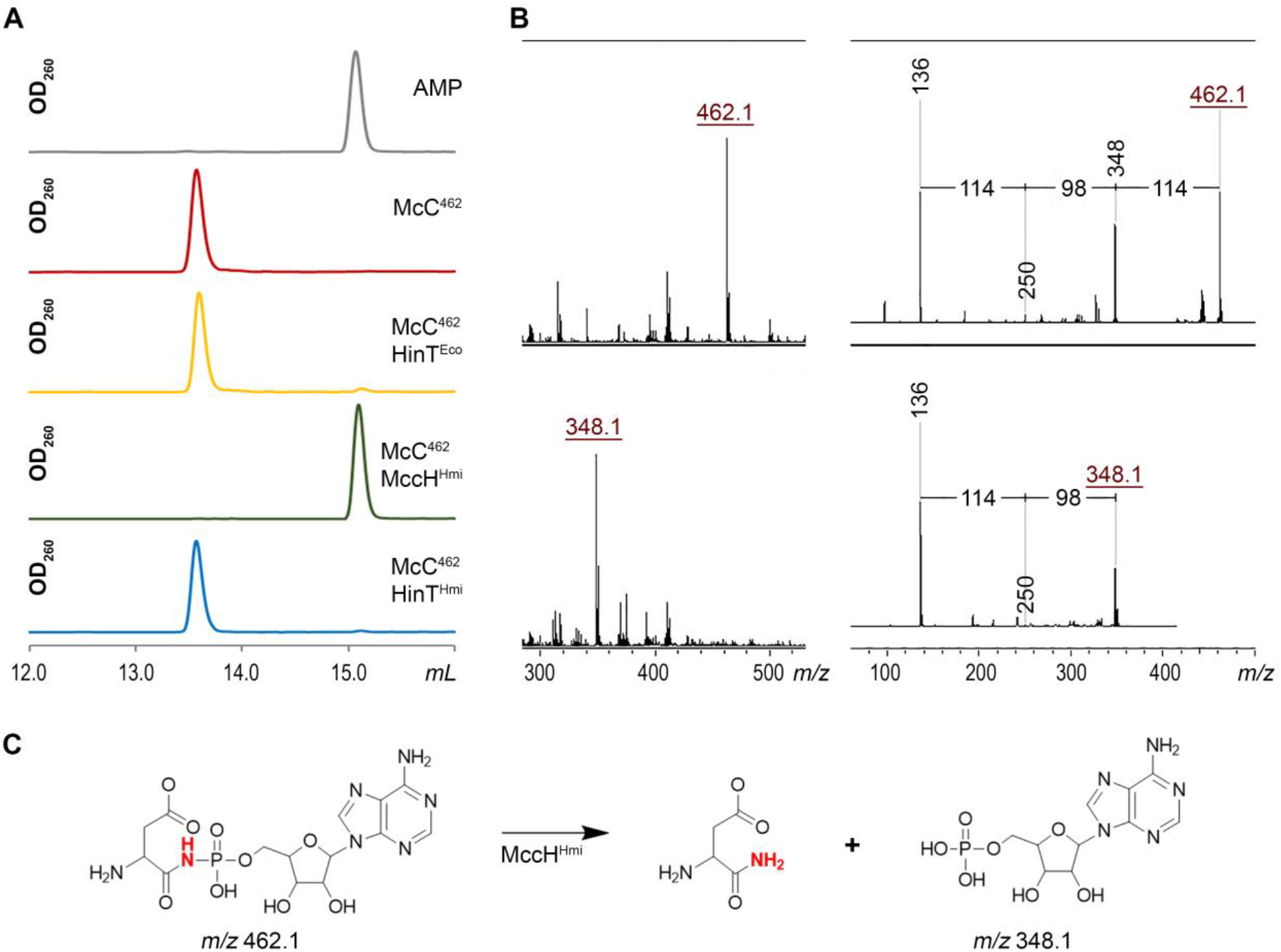
MccH^Hmi^ hydrolyzes aspartamide-adenylate McC^462^. (A) RP HPLC elution profiles of the products of incubation of processed aspartamide-adenylate McC^462^ with MccH^Hmi^, HinT^Hmi^, and HinT^Eco^. (B) MALDI-TOF-MS and MS-MS fragmentation analyses of McC^462^ and the product of its hydrolysis by MccH^Hmi^. For description of mass-ions, refer to the text. (C) A scheme of aspartamide-adenylate hydrolysis reaction by MccH^Hmi^.

In the presence of Hint^Eco^ or HinT^Hmi^, McC^462^ remained largely intact, with only trace amounts of AMP formed in the course of the reaction. We, therefore, decided to assess whether HinT^Hmi^ is an active phosphoramidase using AMP-N-ε-(N-α-acetyl-lysine methyl ester)-5ʹ-phosphoramidate (εK-AMP), a previously described HinT phosphoramidase model substrate (17). Incubation of εK-AMP with HinT^Hmi^, HinT^Eco^, or MccH^Hmi^, followed by RP-HPLC and MALDI-TOF-MS, revealed that both HinT^Hmi^ and HinT^Eco^ hydrolyzed it with the release of AMP, while MccH^Hmi^ did not (Figs. 5A-C). We, therefore, conclude that the MccH^Hmi^ hydrolase cleaves the P-N bond in aspartamide-adenylate but not in εK-AMP. HinT^Hmi^ has a different specificity - it is largely inactive towards processed McC^1120^ but hydrolyzes εK-AMP well. HinT^Hmi^ and HinT^Eco^ have different specificity – they hydrolyze εK-AMP well but are largely inactive towards processed McC^1120^ (Fig. 4A and 5A). These results explain why both HinT enzymes failed to provide resistance to McC-in the antibiotic susceptibility test at our conditions in vivo (Fig. 3).

**Figure 5.**
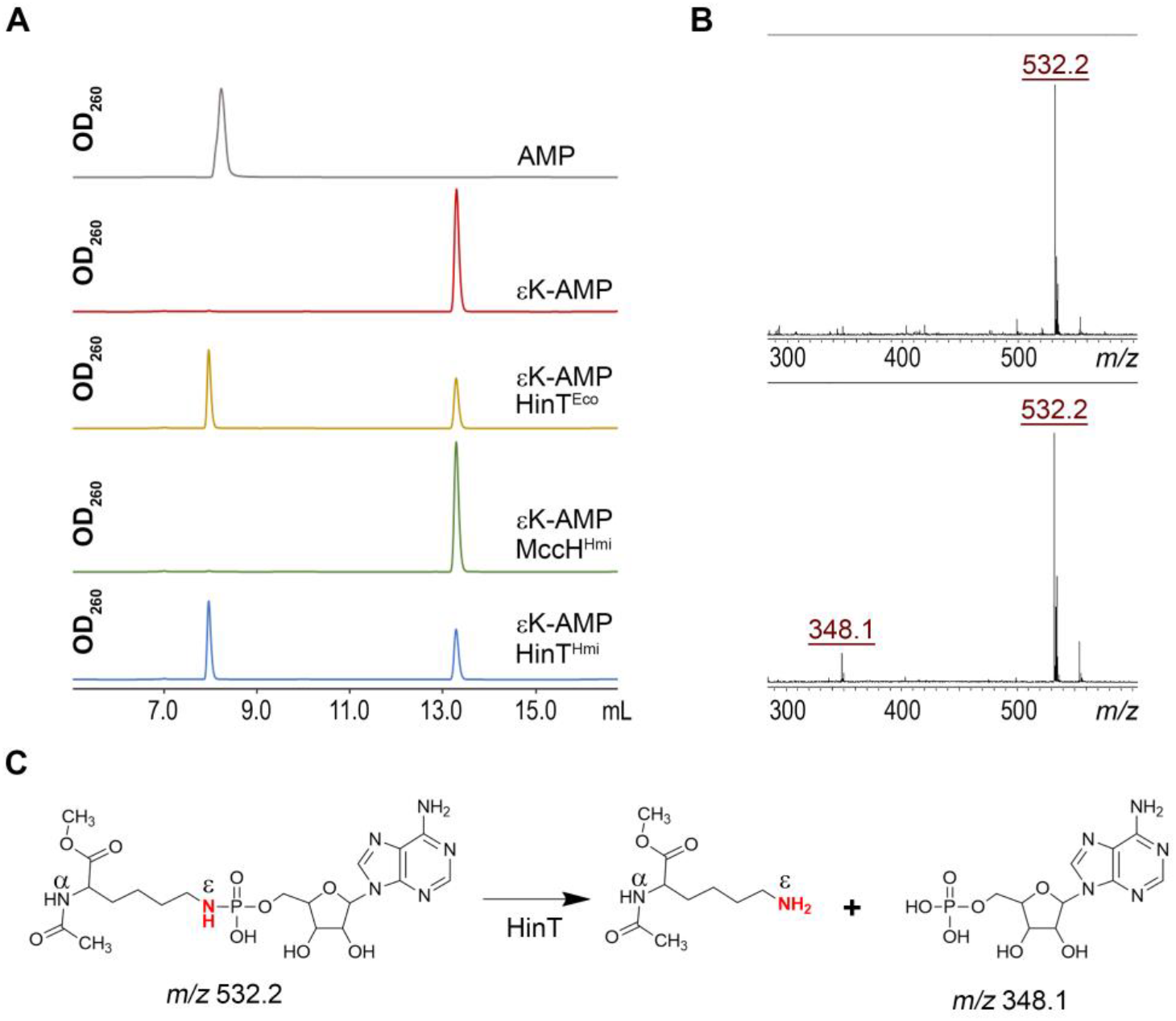
Substrate specificity of MccH^Hmi^, HinT^Hmi^, and HinT^Eco^ phosphoramidases. (A) HPLC elution profiles of products of incubation of εK-AMP with HinT^Eco^, HinT^Hmi^, or MccH^Hmi^. (B) MALDI-TOF-MS analysis of the hydrolysis reaction of εK-AMP by HinT^Hmi^ protein. (C) The scheme of HinT-mediated hydrolysis reaction of εK-AMP (36).

### MccH homologs are present in diverse bacteria

HIT domain-containing proteins are widespread among prokaryotes (see Methods for details on domain identification). A phylogenetic tree constructed using available HIT domain sequences revealed that MccH^Hmi^ belongs to a distinct clade highlighted by red color in Fig. 6A (see also Fig. S4). This clade also contains proteins from putative *mcc*-like clusters from *Nocardiopsis*, *Pseudomonas*, and *Thermobifida*, as well as multiple proteins encoded by standalone genes.Genes encoding MccH homologs from the *mcc*-like cluster from *Nocardiopsis kunsanensis* DSM44524, as well as standalone genes from *Pseudomonas fluorescens* A506, *Salmonella enterica* ser. Newport, *Microcystis aeruginosa* PCC9809, and *Parcubacteria* strains GWA24037 and DG742 were cloned into *E. coli* expression vector, and the ability of the resulting plasmids to make *E. coli* cells resistant to the two forms of McC was tested. As can be seen from Fig. 6B, overexpression of most MccH-like genes led to resistance to McC^1120^ but not to McC^1177^. The apparently inactive MccH-like proteins from *M. aeruginosa* strain PCC9809 and *Parcubacteria* DG742 are the earliest branching MccH homologs tested, and they may have evolved different substrate specificity.

**Figure 6.**
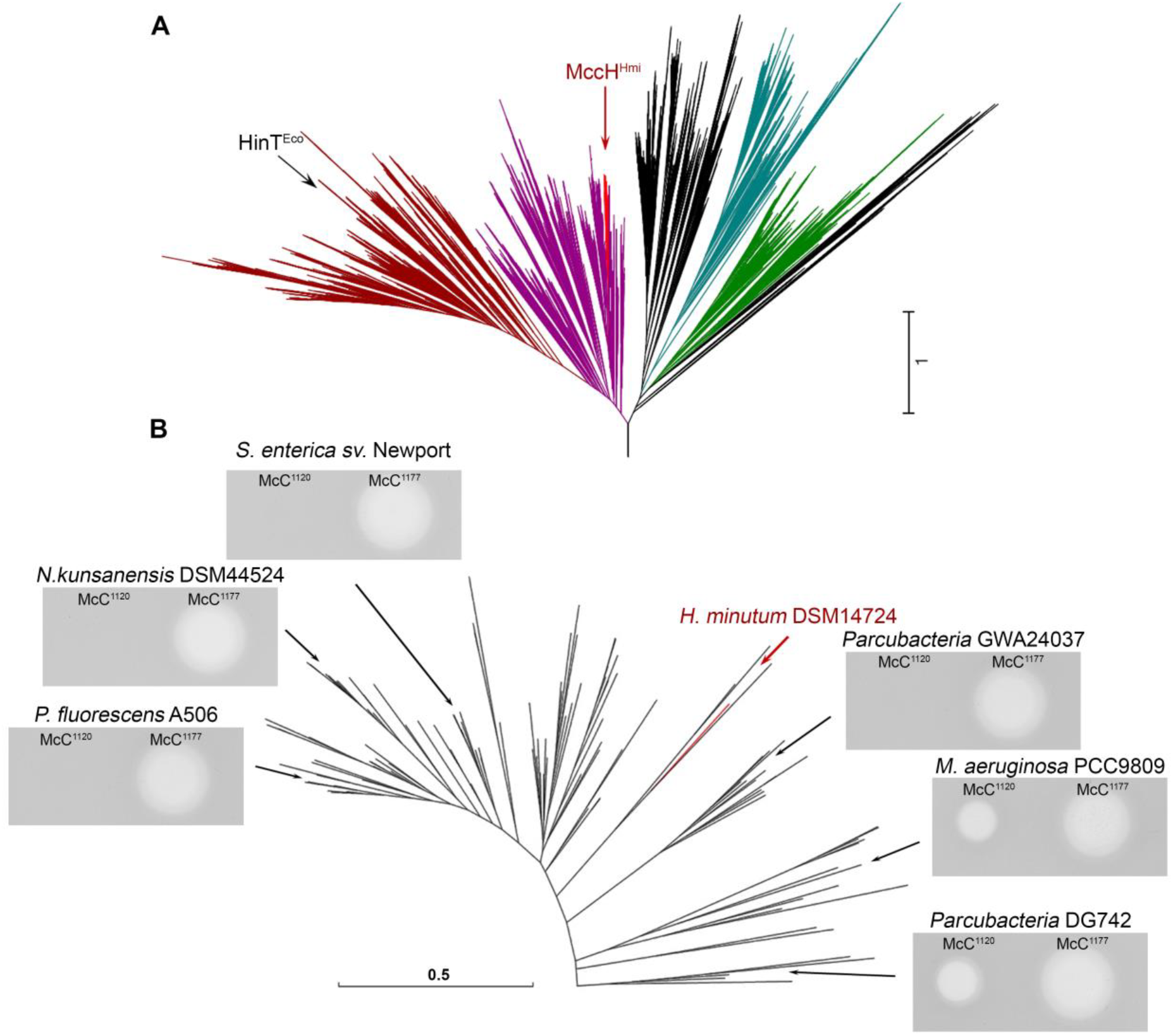
Diversity of McC – specific phosphoramidases. (A) The approximate maximum likelihood (ML) phylogenetic tree of the HIT domain proteins from completely sequenced genomes. Dark magenta and dark red - HinT-like proteins (dark red shows the Protein Kinase C Interacting protein related subgroup); bright red - the MccH^Hmi^ clade that is expanded in (B); cyan - GalT; green - FHIT. In black are clades without any clear profile signature. The arrows point to HinT^Eco^ (within the PKCI clade) and MccH^Hmi^. (B) The approximate ML phylogenetic tree of proteins within the MccH^Hmi^ clade and growth inhibition zones formed by McC^1120^ and McC^1177^ on lawns of *E. coli* B cells transformed with plasmids expressing MccH^Hmi^ or homologs from indicated positions on the tree. The MccH^Hmi^ branch is highlighted in red.

### Mutational analysis of MccH active center

The mechanism of nucleotide phosphoramidate hydrolysis is best studied for the *E. coli* enzyme, HinT^Eco^ (19), and its human homologue, hHint1 (20, 21). The characteristic feature of the HIT superfamily proteins is a conserved histidine triad motif, H*x*H*x*H*xx*, where H is a histidine, and *x* are hydrophobic residues (12). The three essential catalytic histidines form a network of hydrogen bonds with the substrate that promote proton transfer from protonated C-terminal histidine of the triad (H103 in HinT^Eco^ or H102 in HinT^Hmi^) to phosphoramidate unbridged oxygens and amide nitrogen and facilitate nucleophilic attack of the central histidine (HinT^Eco^ H101 or HinT^Hmi^ H100) on the phosphorus atom resulting in P-N bond hydrolysis (19, 21, 22). Another conserved His residue, H39 in HinT^Eco^ (H38 in HinT^Hmi^) located outside of the triad motif closer to the N-terminus of HIT enzymes, contributes to catalysis by stimulating the protonation of the third histidine of the triad by stabilizing its cationic state (21). Mutational analysis of human Hint1, a close homologue of Hint^Eco^, revealed that substitutions of conserved histidines equivalent to H39 and H103 in HinT^Eco^ substantially reduce the catalytic activity (22).

Interestingly, HIT proteins of the MccH clade contain a modified motif where the third histidine is substituted for lysine (K103 in MccH^Hmi^). Together with this substitution, MccH-like proteins lost the N-terminal conserved H residue, which is replaced by phenylalanine (F44 in MccH^Hmi^) (Fig. 7A, Supplementary Fig. S4). Structural modeling (see below) indicates that F44 makes hydrophobic contacts with the side chain of K103 in MccH^Hmi^ (Fig. 8A). In addition, all members of MccH clade acquired glutamate (E93 in MccH^Hmi^) five residues away from the triad motif. In the structural model of MccH^Hmi^, E93 makes a hydrogen bond (or a salt bridge) with K103, thus playing the same functional role as H39 in HinT^Eco^ (H38 in HinT^Hmi^) (Fig. 8). Since this triple substitution should still allow phosphoramide bond hydrolysis, we speculate that the positively charged lysine occupying position of H103 in MccH^Hmi^ ultimately donates its stationary proton to the nitrogen of the phosphoramide bond (Fig. 8B). To test this conjecture and better understand the origin of MccH-like proteins specificity, we prepared two MccH^Hmi^ mutants harboring single substitutions K103H and F44H. A double MccH^Hmi^ K103H/F44H mutant was also engineered. As a key residue in the triad, H101 in MccH^Hmi^ is supposed to directly participate in catalysis. Therefore, a protein with substitution H101N was prepared and tested for phosphoramidase activity. As expected, the H101N mutant, which served as a control, was catalytically inactive, *i.e*., no hydrolysis of processed McC^1120^ was detected (Fig. 7B). The K103H substitution had also eliminated the hydrolytic activity of MccH^Hmi^, confirming that a lysine characteristic for MccH-like proteins is essential for the catalytic function. The F44H mutant retained its activity, which is also an expected result. The double K103H/F44H mutant was insoluble, and thus its enzymatic properties could not have been assessed. To confirm the *in vitro* hydrolytic activity of the mutants, the *E. coli* cells harboring the corresponding plasmids were tested for their susceptibility to McC^1120^. As shown in Fig. 7C, H101N, and K103H substitution completely abolished immunity to peptidyl-adenylates, while the phenotype of MccH^Hmi^ F44H-expressing cells was indistinguishable from that of the cells producing wild type MccH^Hmi^.

**Figure 7.**
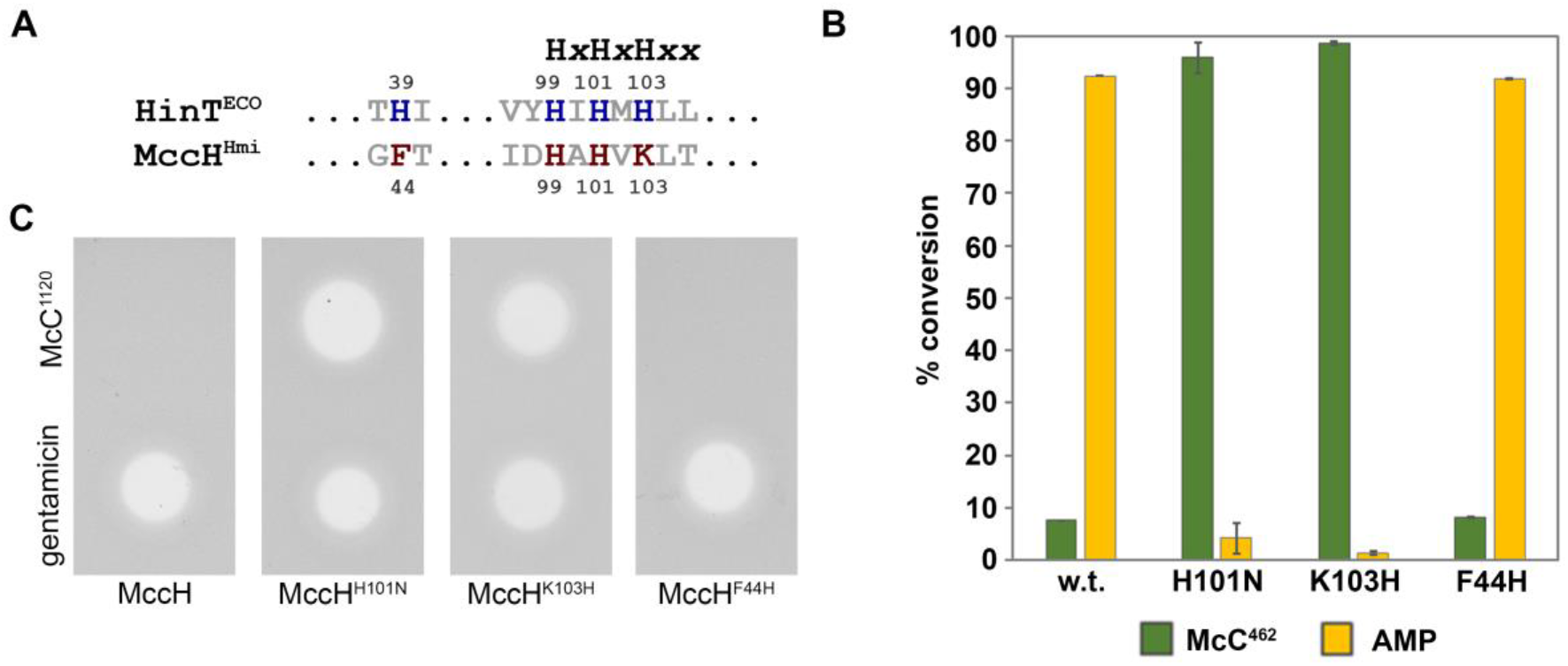
Mutational analysis of active site residues of MccH^Hmi^. (A) Sequence alignment of the conserved HIT motif of HinT^Eco^ and MccH^Hmi^. (B) Phosphoramidase activity of MccH^Hmi^ active site mutants. *In vitro* reactions containing aspartamide-adenylate McC^462^ were incubated with wild-type MccH^Hmi^ and the mutant proteins, containing H101N or K103N substitution, then analyzed by RP-HPLC. The conversion of McC^462^ was calculated as the percentage of McC^462^ and AMP absorption peak areas remained after the reaction completion relative to corresponding peak areas observed without the addition of enzymes. The bars represent the mean conversion percentage values calculated from three independent measurements ± SD. (C) Mutations in the active center of MccH^Hmi^ abolish immunity to McC^1120^. Growth inhibition of *E. coli* cells harboring pBAD plasmids encoding indicated proteins. 5 μM solutions of McC^1120^ and 0.5 μg/mL of gentamicin were deposited on freshly prepared lawns and allowed to grow overnight at 37 °C in conditions of induction of plasmid-borne genes.

**Figure 8.**
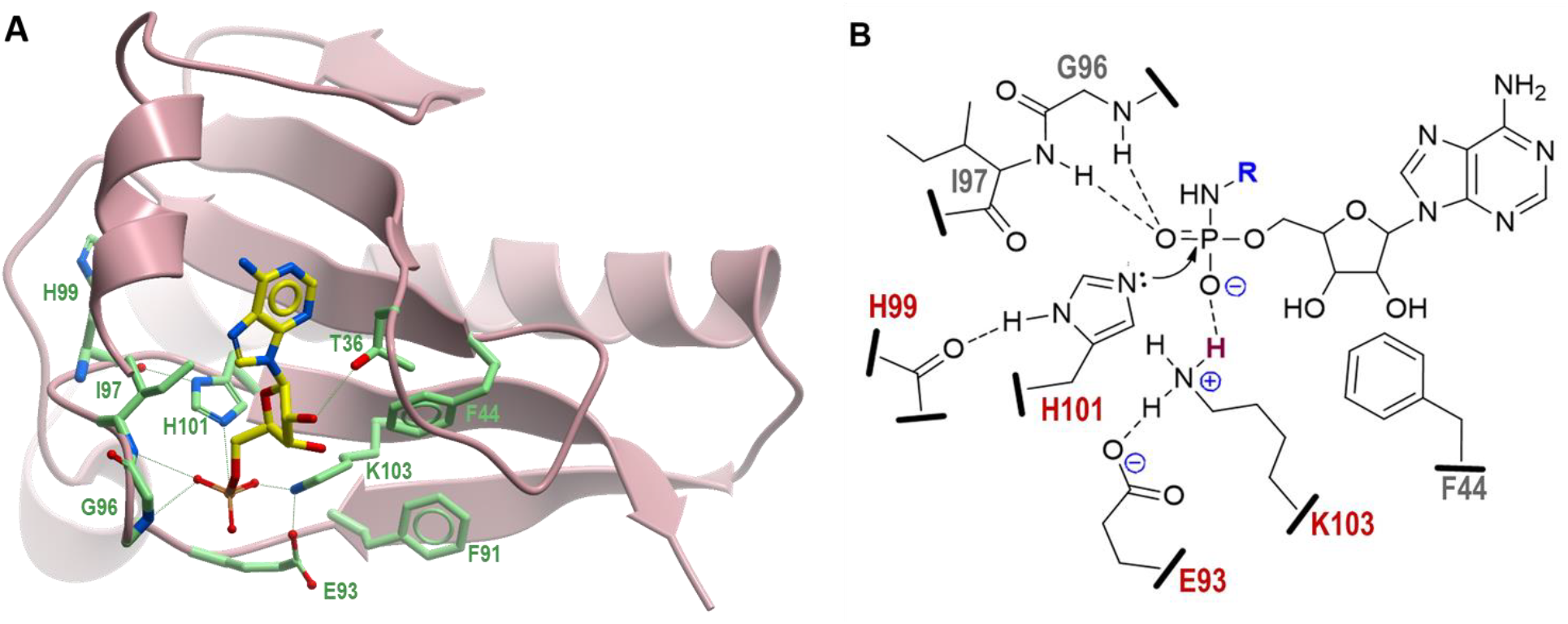
3D-structural model of MccH^Hmi^ and proposed catalytic mechanism. (A) Model structure of MccH^Hmi^ generated by SWISS-MODEL homology modeling server (23) using as template the crystal structures of HIT-like protein from *Mycobacterium paratuberculosis* (PDB:3P0T) (24) and human HINT1-AMP complex (PDB:3TW2) (28). Positions of catalytic residues in the MccH^Hmi^ active site in complex with AMP are shown as CPK colored sticks. C-atoms in AMP are in yellow, C-atoms in side chains are in light green. (B) Schematic diagram representing the active site of MccH^Hmi^ with the bound substrate, aspartamide-adenylate (aspartyl moiety is represented by R and shown in blue). Residues of the E*xxxxx*H*x*H*x*K*xx* motif conserved among all members of the MccH clade that are directly involved in the catalysis are shown in red. Potential hydrogen bonds between peptide side chains, backbone atoms, and phosphate oxygens are indicated by dashed lines. Proton to be transferred from the ε-amino group of K103 to unbridged oxygen of the phosphoramide is depicted in dark red. Nucleophilic attack of the unprotonated nitrogen of H101 on electrophilic phosphorus atom is shown by a curved arrow.

### Structural model of MccH^Hmi^

The loss of functional activity by MccH^Hmi^ mutant K103H suggested that this residue, specific to MccH clade proteins, is involved in substrate binding and/or catalysis. We also hypothesized that some other active site residues might spatially constrain the catalytic pocket environment favoring the flexible aliphatic side chain of lysine over a more rigid imidazole ring of histidine. To explore the possible spatial organization of the active center of MccH^Hmi^, we generated its 3D-model using the SWISS-MODEL homology modeling program (23). The resulting model of MccH^Hmi^ (Fig. 8A) is based on a top-ranked *Mycobacterium paratuberculosis* HIT-like protein structure (PDB:3P0T) (24) with a GMQE quality score of 0.68 indicative of good reliability and accuracy. To reveal the potential interactions in the substrate-binding pocket, the model structure of MccH^Hmi^ was superimposed with the crystal structure of the human histidine triad nucleotide-binding protein 1 (hHINT1) in complex with AMP (PDB: 3TW2) (25).

As expected, the overall structure of MccH^Hmi^ and the spatial organization of its active site are very similar to that of other HinT and HIT-like proteins, with a notable exception of the C-terminal non-conserved 45 amino acids that model differently depending on the homology template used. In the model, MccH^Hmi^ forms a symmetric homodimer with each protein monomer capable of binding and hydrolyzing the substrate (Supplementary Fig. S5). The nucleoside-binding pocket is formed mostly by conserved hydrophobic residues F11, F12, L15, F34, P37, V46, F38, and I97. The hydroxyl group of T36 makes a hydrogen bond with ribose 2’-OH in AMP, thus contributing to nucleotide recognition. The N-atoms of the side chains of catalytic H101 and K103 are positioned (2.5-2.7Å) to make strong hydrogen bonds with the P- and unbridged O-atoms of the phosphate moiety of AMP, respectively, (Fig. 8A). The interatomic distances (2.9-3.6Å) between the peptide backbone amide and carbonyl groups of residues G96, I97, and H99 and their interacting partners (unbridged oxygen, and protonated N-atom of H101, respectively) are within a range optimal for hydrogen bonding consistent with the proposed catalytic mechanism (Fig. 8B). Unexpectedly, the carbonyl group of E93, a conserved residue among members of the MccH clade, is in close proximity (2.3Å) to the protonated N-atom of K103, suggesting a strong hydrogen bond or salt bridge that would stabilize the charged state of K103 and facilitate the catalysis. Furthermore, consistent with the results of our mutagenesis experiments (Fig. 7), the bulky hydrophobic side chains of F44 and F91 replacing conserved H39 and I86 in HinT^Eco^ and HinT^Hmi^, about the side chain of K103, stabilize its conformation and direct it towards the phosphate. Thus, the positioning of F44, F91, and E93 in the active center explain the observed preference for catalytic lysine in MccH instead of histidine in HinT clades.

## DISCUSSION

In this work, we uncover a novel mechanism of immunity to microcin C-like compounds by MccH^Hmi^, a HIT-like phosphoramidase encoded in the *mcc* cluster of *H. minutum*. The cluster produces two separate peptide-adenylates that are analogous to McC^1120^, a toxic maturation intermediate of *E. coli* McC that lacks the aminopropyl decoration. As of today, the *mcc-*operon from *H. minutum* is the only validated operon that produces two McC-like compounds with different peptide parts. Since peptide parts determine the specificity of antibacterial action by allowing selective import into sensitive cells (26), *H. minutum* DSM 14724 may target distinct, non-overlapping sets of its competitors by McC-like compounds it produces.

Like other McC-producing organisms, *H. minutum* should experience the buildup of toxic processed products inside the cell, which could lead to cessation of protein biosynthesis. The MccH^Hmi^ enzyme, the product of the *mcc* operon, alleviates this problem by cleaving the bond between phosphorus and nitrogen in the toxic aspartamide-adenylate that is produced after proteolytic processing of either of the two McC-like compounds encoded by the operon, thus providing self-immunity to the producing cell. MccH^Hmi^ makes cells resistant to the maturation intermediate of *E. coli* McC that lacks the aminopropyl decoration but not to fully mature McC^1177^. The catalytic mechanism of phosphoramide hydrolysis requires a transient protonation of two unbridged oxygens (21, 22). The presence of an additional aminopropyl group on the phosphate in unprocessed McC^1177^ and processed McC^519^ precludes the proton transfer reaction and renders the phosphorus center inaccessible to nucleophilic attack by the catalytic histidine (HinT^Eco^ H101, HinT^Hmi^ H100, or MccH^Hmi^ H101). Thus, *H. minutum* and *E. coli mcc* operons use different strategies to overcome the self-intoxication of producers. The *H. minutum mcc*-operon produces two peptidyl-adenylates without additional modifications, and the processing of both compounds leads to identical toxic aspartamide-adenylate. MccH^Hmi^ hydrolyzes the phosphoramide bond in aspartamide-adenylate with the formation of AMP and aspartamide. The absence of additional genes in the *H. minutum mcc-*operon that may be involved in self-immunity suggests that MccH^Hmi^ is sufficient to counter the inhibitory effects caused by the buildup of the toxic product. In *E. coli*, the MccD/E enzyme complex installs the aminopropyl decoration at the phosphate of peptide-adenylate, which allows increasing the potency of antibacterial action ~10-fold by increasing the affinity of the processed compound to its target, AspRS (4). The presence of activity-enhancing decoration renders the MccH^Hmi^ enzyme inactive, necessitating another mechanism to overcome self-intoxication. MccE detoxifies both aminopropylated and non-aminopropylated aspartamide-adenylates by acetylating the amino group of the aspartate (8).

The structural model of MccH^Hmi^ built based on a crystal structure of homologous HIT-like protein from *M. paratuberculosis* (PDB:3P0T) (24) provides a plausible view on a spatial organization of the active center and offers clues to understanding the enzyme’s substrate specificity. Importantly, the model points to the functional role of the conserved hydrophobic (F44, F91) and charged (E93) residues in the activation of catalytic K103 for hydrolysis of aspartamide-adenylate that can be tested experimentally. It is also predicted that residues M95 and W115 of the C-terminal loop of one MccH monomer together with L110 from the adjacent C-terminal loop of the other MccH monomer form a tight aspartamide-binding site which would sterically occlude the binding of bulkier groups such as ε-lysineamide of εK-AMP. This view is consistent with the fact that HinT^Eco^ lacking the C-terminal extension present in MccH^Hmi^ was active towards εK-AMP, but could not hydrolyze aspartamide-adenylate.. It is also supported by previous observations that both deletion and swapping of the C-terminal loop between human HINT1 and HinT^Eco^ strongly affect both the catalytic activity and substrate specificity (19, 27, 28). The proposed model will be validated in our future genetic, biochemical, and structural studies of bacterial MccH and HinT proteins.

Previous studies have shown that bacterial and human HinTs exhibit a broad substrate specificity; they can accommodate both purine and pyrimidine nucleotides with various substitutions in the aminoacyl moiety, including D- and L-stereomers of tryptophane and sterically hindering N-ε-(N-α-acetyl-lysine methyl ester)-adenosine phosphoramidates (11, 27). Unlike HinT, the MccH^Hmi^ and its homologues from four diverse bacterial species characterized in this work apparently have evolved much more specialized enzymes that show a clear preference for aspartamide-adenylate (Figs. 4 and 5). Our results suggest that the fully processed McC is a *bona fide* substrate for MccH. We speculate that MccH-like proteins from *M. aeruginosa* PCC9809 and *Parcubacteria* DG742, which are inactive towards *E. coli* McC, may recognize yet unidentified species-specific McC-like compounds that carry different nucleoside or amino acid moieties.

## MATERIALS AND METHODS

### Molecular cloning

*E. coli* DH5α was used for cloning. All primers were synthesized by Evrogen (Russia); their sequences are listed in Supplemental Table S1. The *H. minutum* DSM 14724 or *E. coli* BW25113 genomic DNA was used as a template for PCR. Phusion DNA polymerase (Thermo Scientific) was used for PCR.

For MccB_1_^Hmi^ and MccB_2_^Hmi^ activity analysis, their coding sequences were amplified from the *H. minutum* DSM 14724 genomic DNA. The PCR products were digested with BamHI and SalI restriction endonucleases and inserted under the same sites into the pET22_MBP vector (26) to create an N-terminal fusion protein with MBP-tag.

For the heterologous *mcc*^*Hmi*^ expression system, DNA fragment spanning *mccP1,P2,P3*^*Hmi*^ genes was amplified from genomic DNA then digested with EcoRI and KpnI and inserted into pACYCDuet-1 vector (Novagen-Millipore, USA) linearized with the same restriction endonucleases. To construct MccA-MccB expression vectors, first, the phosphorylated self-complementary oligonucleotides containing sequences of MccA_1_^Hmi^ and MccA_2_^Hmi^ ORFs were inserted into pRSFDuet-1 vector (Novagen-Millipore, USA), digested with NcoI and HindIII, resulting in pRSF-MccA_1_^Hmi^ and pRSF-MccA_2_^Hmi^ vectors, respectively. Then, *mccB*_*1*_^*Hmi*^ and *mccB*_*2*_^*Hmi*^ were PCR-amplified, digested with NdeI and KpnI, and inserted into pRSF_*mccA*_*1*_^*Hmi*^ and pRSF_*mccA*_*2*_^*Hmi*^ vectors linearized with NdeI and KpnI resulting in pRSF_*mccA*_*1*_*B*_*1*_^*Hmi*^ and pRSF_*mccA*_*2*_*B*_*2*_^*Hmi*^, respectively.

To obtain an arabinose inducible vector for heterologous expression of HIT proteins, the pBAD/His B vector (Invitrogen-Thermo Fisher, USA) was linearized by PCR with appropriate primers that contained an introduced ribosomal binding site and SalI and HindIII restriction sites to generate pBAD30_SalRBS. Next, the *mccH*^*Hmi*^, *hinT*^*Hmi*^, and *hinT*^*Eco*^ genes were PCR-amplified using genomic DNA as a template and correspondent primers. Genes encoding the homologs of MccH^Hmi^ were purchased as synthetic DNA fragments from IDT, USA. All amplified PCR products and synthetic fragments were digested with SalI and HindIII and inserted between the same sites into the pBAD30 expression vector.

To create vectors for HIT protein fused with C-terminal His6 tags for protein purification, *mccH*^*Hmi*^, *hinT*^*Hmi*^, and *hinT*^*Eco*^ genes were PCR-amplified, digested with NdeI and XhoI and inserted into pET22(b) vector (Novagen-Millipore, USA). The site-directed mutagenesis of *mccH*^*Hmi*^ was carried out using overlap extension PCR (29) with appropriate primers.

### Recombinant protein expression and purification

Recombinant proteins were produced in *E. coli* BL21DE3 transformed with the appropriate plasmid. The cells were grown in 500 mL of TB medium, supplemented with ampicillin, to an OD_600_~0.7 and induced with 0.2 mM isopropyl-β-D-thiogalactopyranoside (IPTG). After induction, the culture was transferred for overnight growth at 18 °C, 180 rpm. The cells were harvested by centrifugation at 8 000 x *g* at 4 °C for 20 min, resuspended in ice-cold resuspension buffer (20 mM Tris-HCl, 300 mM NaCl, 1mM DTT, pH 8.0), supplemented with 1 mM PMSF, and disrupted by sonication. For the His-tagged proteins, imidazole was added up to 2 mM. The lysate was cleared by centrifugation at 30 000 x *g* at 4 °C for 20 minutes. The cleared lysate was applied to a pre-equilibrated column with a tag-binding resin; depending on the tag present, either Amylose Resin (NEB) or Talon CellThru Co^2+^-chelating resin (Takara-Clontech) were used. The resin was washed with 10 column volumes of resuspension buffer, followed by elution with 5 column volumes of the Elution Buffer (20 mM Tris-HCl, pH 8.0, 50 mM NaCl, 10% glycerol) supplemented with either 10 mM maltose or 0.5 M of imidazole.

### Adenylation of MccA_1_^Hmi^ and MccA_2_^Hmi^

For validation of MccA/MccB pairs of *H. minutum mcc*-like gene cluster, the *in vitro* modification of synthetic MccA_1_^Hmi^ (MNDKATIEIKKDEKKAEPKKVVVVKTSIKAGPAAFN) and MccA_2_^Hmi^ (MNEKTAQESQKTESPKAETPAKKAVIVKTRIKAGPGGGGLVHPIAN) (GeneScript, USA) peptides using purified MccB_1_^Hmi^ and MccB_2_^Hmi^ reactions was performed. Reaction mixtures contained 50 μM of synthetic peptides (either MccA_1_^Hmi^ or MccA_2_^Hmi^), 5 μM recombinant MccB_1_^Hmi^ or MccB_2_^Hmi^ and 2 mM of each NTP in the reaction buffer [50 mM Tris-HCl pH 8.0, 150 mM NaCl, 10 mM MgCl_2_, 5 mM DTT]. The reaction was carried out at 30 °С for 16 h, then stopped by the addition of 0.1% trifluoracetic acid (TFA) in water. The products of the reaction were analyzed by MALDI-TOF-MS for the presence of adenylated MccA_1_^Hmi^ and MccA_2_^Hmi^.

### McC_1_^Hmi^ and McC_2_^Hmi^ production test

*E. coli* BL21 (DE3) cells harboring a combination of either pRSF_*mccA*_*1*_*B*_*1*_^*Hmi*^ and pACYC_*mccP*_*1*_*P*_*2*_*P*_*3*_^*Hmi*^ or pRSF_*mccA*_*2*_*B*_*2*_^*Hmi*^ and pACYC_*mccP*_*1*_*P*_*2*_*P*_*1*_^*Hmi*^ plasmids were grown in 25mL of 2YT medium at 30 °C with constant shaking at 180 rpm. Upon reaching OD_260_ ~0.7, the cells were induced with 0.25 mM IPTG and grown for an additional 20 h at 30 °C. After that, the cells were pelleted at 5 000 x *g* and resuspended in 250 μL of M9 medium. Aliquots of 15 μL were deposited on a freshly prepared M9 agar lawn of *E. coli* strain B harboring pRSFduet-1, pETduet-1, and pACYCduet-1 vectors. Plates were incubated at 30 °C overnight to allow the formation of the lawn. The next day the plates were inspected for the presence of growth inhibition zones. Additionally, the cells and the surrounding agar were analyzed for the presence of McC-like products using mass spectrometry.

### *In vivo* immunity assay

*E. coli* strain B was transformed with pBAD_SalRBS vectors containing the *mccH*^*Hmi*^ gene or its homologues. The overnight cultures of the *E. coli* cells with the appropriate vectors were diluted 1000-fold in M9 agar medium, supplemented with 1% glycerol, 0.02% yeast extract, 10 mM arabinose, and 100 μg/mL ampicillin. The sensitivity of cells carrying plasmid-borne genes encoding HIT-like proteins was measured by applying 5 μL drops of 5 μM McC^1120^ and 5 μM McC^1177^ on the surface of the plate and allowed to dry, gentamycin (0.5 μg/mL) was used as a control growth inhibitor. Plates were incubated for 16 h at 30 °C for a lawn to form. The next day the growth inhibition zones were analyzed.

### AMP-N-ε-(N-α-acetyl lysine methyl ester) 5ʹ-phosphoramidate (εK-AMP) synthesis

The synthesis of εK-AMP was performed as described elsewhere with minor modifications (17). To synthesize εK-AMP, 0.12 g (6.25 mmol) of 1-ethyl-3-(3-dimethyllaminopropyl) carbodiimide HCl (Sigma Aldrich) was added to the flask containing 0.98 g (2.7 mmol) of AMP (Sigma Aldrich), 0.1 g of (0.42 mmol) N-α-acetyl-L-lysine methyl ester HCl (Sigma Aldrich) and 10 mL of ultrapure water, the pH of the solution was adjusted with triethylamine to a value of 7.5. The mixture was incubated in the shaker at 65 °C, with vigorous shaking at 250 rpm for 22 h. After allowing the reaction mix to cool down to room temperature, the mixture was lyophilized, then re-dissolved in 0.1% TFA in water and applied to Luna 5 µm C18 100 Å, LC Column (250×10 mm) (Phenomex). The purification of εK-AMP was performed in a linear (5–25%) gradient of acetonitrile. The fractions were analyzed for the presence of εK-AMP by MALDI-TOF mass spectrometry, fractions containing εK-AMP were subjected for additional chromatographic purification on the same column in a linear gradient of acetonitrile (0-25%) in triethylammonium acetate (TEAA) buffer, pH 6.5.

### *In vitro* phosphoramidase activity assay

To test the phosphoramidase activity of MccH^Hmi^, HinT^Hmi^, HinT^Eco^ enzymes, and their respective variants, the processed forms of *E. coli* McC^1177^ and McC^1120^ – (McC^519^ and McC^462^, respectively) and εK-AMP were used as substrates. For production, purification, and processing of McC forms, refer to Metlitskaya *et al.* (6). 50 μM of processed McC or εK-AMP were mixed with 5 μM of the enzyme in the reaction buffer [20 mM HEPES pH 7.2, 2.5 mM MgCl_2_, 2.5 mM MnCl_2_]. The reaction was incubated at 25 °С for 30 minutes, terminated by the addition of 0.1% TFA and analyzed for hydrolysis by HPLC and mass spectrometry.

### Reverse Phase HPLC analysis of the products of the *in vitro* reactions

All biochemical reactions were analyzed on 1220 Infinity II LC System (Agilent), the peaks separation occurred on Zorbax Eclipse Plus C18 5 µm (4.6×250 mm) column (Agilent) in 0.1 M TEAA buffer system, pH 6.0 in the varying linear gradient of acetonitrile.

The products of McC^462^ hydrolysis reactions were separated in a linear gradient of acetonitrile (0-20%) over a period of 15 minutes. After incubation of McC^519^ with HIT enzymes, the reaction products were separated in the acetonitrile gradient (0-22%) for 15 minutes. After hydrolysis of εK-AMP by HIT enzymes, the reaction products were analyzed in the linear acetonitrile gradient (5-30%) lasting for 15 minutes. The chromatograms were processed with the use of ChemStation software (Agilent) and elution profiles were exported in comma-separated values format.

### Mass spectrometry analysis

1-2 μl of the sample aliquots were mixed with 0.5 μl of matrix mix (Sigma Aldrich) on a steel target. The mass spectra were recorded on a mass spectrometer UltrafleXtreme MALDI-TOF/TOF (Bruker Daltonics) equipped with a neodymium laser. The molecular MH^+^ ions were measured in reflector mode; the accuracy of measured results was within 0.1 Da.

### Sequence analysis

Proteins, containing the histidine catalytic triad were identified in 4,621 completely sequenced genomes available in 2016. Profiles, belonging to the NCBI CDD (30) superfamily cl00228 were used as PSI-BLAST (31) queries to search the protein sequences, encoded in this set. The resulting set of 10,580 proteins was clustered using UCLUST (32) and aligned using MUSCLE (33) (Supplementary Fig. S6); alignments were iteratively compared to each other using HHSEARCH and aligned using HHALIGN programs (34). The approximate ML tree was reconstructed using the FastTree program (35) with WAG evolutionary model and gamma-distributed site rates.

## Acknowledgements

This work was supported by NIH RO1 grant AI117270 (to KS and Satish A. Nair) and Skoltech institutional funds to KS, the Russian Science Foundation grants RSF #16-14-10356 and RSF #19-14-00266 to SD, Rowan University departmental funds to SB. Y.I.W. is supported through the intramural program of the U.S. National Institutes of Health. Bioinformatic analysis was partially supported by the Ministry of Science and Higher Education of the Russian Federation grant 075-15-2019-1661. MALDI-TOF-MS facility became available to us in the framework of the Moscow State University Development Program PNG 5.13.

## Supplementary materials

**Figure S1. Production of McC_1_^Hmi^ and McC_2_^Hmi^ in heterologous host.**

(A) *E. coli* cells harboring plasmid-borne *H. minutum mcc* operon do not produce toxic compounds: *mccA*_*1*_*B*_*1*_ – *E. coli* BL21(DE3) cells harboring pRSF_*mccA*_*1*_*B*_*1*_^*Hmi*^ and pACYC_*mccP*_*1*_*P*_*2*_*P*_*3*_^*Hmi*^ plasmids, *mccA*_*2*_*B*_*2*_ – BL21(DE3) cells carrying pRSF_*mccA*_*2*_*B*_*2*_^*Hmi*^ and pACYC_*mccP*_*1*_*P*_*2*_*P*_*3*_^*Hmi*^ plasmids, control - *E. coli* BL21(DE3) cells harboring empty pRSF and pACYC vectors. Cells were induced for 24 h at 30 °С, then extracted as described in (37). 5 μl of 10-times concentrated cell cultures (upper panel) or cellular extractes (lower panel) were deposited on the surface of McC-sensitive *E. coli* B cells lawn (upper panel). 2 μl of 0.5 μg/mL gentamycin solution was used as a control antibiotic.

(B) MALDI-TOF-MS analysis of *E. coli* BL21 cells harboring pRSF_*mccA*_*2*_*B*_*2*_^*Hmi*^ and pACYC_*mccP*_*1*_*P*_*2*_*P*_*3*_^*Hmi*^ (upper panel) and pRSF_*mccA*_*1*_*B*_*1*_^*Hmi*^ and pACYC_*mccP*_*1*_*P*_*2*_*P*_*3*_^*Hmi*^ plasmids (lower panel). At the top spectrum, MH^+^ at *m/z* 5096.6 corresponding to adenylated MccA_2_^Hmi^, peptide-adenylate lacking N-terminal methionine (MH^+^ at *m/z* 4965.6), full-length MccA_2_^Hmi^ precursor peptide (MH^+^ at *m/z* 4767.6), and MccA_2_^Hmi^ lacking N-terminal methionine (MH^+^ at *m/z* 4636.6) are labeled. MH^+^ ions at *m/z* 3637.0 and 4363.5 correspond to *E. coli* proteins. At the bottom spectrum, ions corresponding to adenylated MccA_1_^Hmi^ (MH^+^ at *m/z* 4255.2), peptide-adenylate lacking N-terminal methionine (MH^+^ at *m/z* 4124.2), full-length MccA_1_^Hmi^ precursor peptide (MH^+^ at *m/z* 3926.2), and MccA_1_^Hmi^ lacking N-terminal methionine (MH^+^ at *m/z* 3795.2) are labeled.

**Figure S2. Coomassie-stained SDS polyacrylamide gel showing purified proteins used in the study.** L – PageRuler Plus Prestained protein ladder; 1 – MccH^Hmi^; 2 – MccH^Hmi^ H101N; 3 – MccH^Hmi^ K103H; 4 – MccH^Hmi^ F44H; 5-HinT^Eco^; 6 -HinT^Hmi^.

**Figure S3. Aminopropyl decoration of aspartamide-adenylate protects the compound from the phosphoramidase activity of MccH^Hmi^, HinT^Eco^, and HinT^Hmi^.**

(A) MALDI-TOF-MS spectra of McC^519^ incubated without the enzyme (top panel) and with HinT^Eco^, MccH^Hmi^, and HinT^Hmi^ (lower panels). The MH^+^ ion at *m/z* 519.2 corresponds to aminopropylated aspartamide-adenylate. No MH^+^ ion at *m/z* 405.2 corresponding to hydrolyzed McC^519^ is observed.

(B) RP-HPLC elution profile of products of incubation of McC^519^, processed aspartamide-adenylate with aminopropyl decoration, without the enzyme and with MccH^Hmi^, HinT^Hmi^, and HinT^Eco^.

**Figure S4. Conservation of the amino acid sequence in the Protein Kinase C Interacting protein-related clade of HIT proteins.** Sequence alignment of HinT clade of HIT proteins (HmiDSM14724a, WP_044187632.1 of Hyalangium minutum DSM 14724; TteBAA798, ACZ41971.1, of Thermobaculum terrenum ATCC BAA-798; EcoNCTC9094, WP_096759427.1 of Escherichia coli NCTC 9094; SteATCC33386, ACZ09064.1 of Sebaldella termitidis ATCC 33386) and MccH clade of HIT proteins (HmiDSM14724, WP_044187428.1 of Hyalangium minutum DSM 14724; PflA506, AFJ55311.1 of Pseudomonas fluorescens A506; NkuDSM44524, WP_017574753.1 of Nocardiopsis kunsanensis DSM 44524; SenNewport, ECU0367860.1 of Salmonella enterica subsp. enterica serovar Newport; ParGWA24037, KKR61370.1 of Parcubacteria bacterium GW2011_GWA2_40_37; MaePCC9809, CCI22782.1 of Microcystis aeruginosa PCC 9809; ParDG742, KPJ57467.1 of Parcubacteria bacterium DG_74_2). Residues conserved in either of the two groups are shown in bold and underlined. Histidine-triad active site region is indicated by a red-shaded box. Conserved and partially conserved hydrophobic and polar residues forming the nucleotide-binding pocket of HIT proteins are indicated by an asterisk (*). Substitutions of the active site residues in MccH clade proteins are indicated by (‡). Red boxes mark residues of “inactive” MaePCC9809 and ParDG742 MccH-like proteins, that differ considerably from the MccH consensus.

**Figure S5. 3D-structural model of MccH^Hmi^ dimer in complex with AMP. The two monomers of MccH are depicted as light green and purple colored ribbons diagrams. Residues of the active site that form the substrate-binding pocket are labeled and shown in a stick representation.**

**Figure S6. Alignment of cluster consensus sequences of the HIT domain proteins from completely sequenced genomes.** The phylogenetic tree, constructed from the multiple alignment of 15,351 HIT sequences (Figure 6A) was split into subtrees at the average depth of 1.5 from the tree tips, producing 292 clusters of 4+ sequences. Consensus sequences were derived for each cluster from the corresponding subsets of the alignment (positions, less than 30% conserved denoted by “x”). Positions corresponding to active site histidine residues are highlighted in yellow. Consensus sequence for the MccH^Hmi^ (CON.109), HinT^Eco^ (CON.19), and HinT^Hmi^ (CON.21) clades are shown in red font.

**Table S1. Primers used in the study.**

## REFERENCES

1. Severinov K, Semenova E, Kazakov A, Kazakov T, Gelfand MS. 2007. Low-molecular-weight post-translationally modified microcins. Mol Microbiol 65:1380–1394.

2. Guijarro JI, González-Pastor JE, Baleux F, San Millán JL, Castilla MA, Rico M, Moreno F, Delepierre M. 1995. Chemical structure and translation inhibition studies of the antibiotic microcin C7. J Biol Chem 270:23520–23532.

3. Roush RF, Nolan EM, Löhr F, Walsh CT. 2008. Maturation of an Escherichia coli ribosomal peptide antibiotic by ATP-consuming N-P bond formation in microcin C7. J Am Chem Soc 130:3603–3609.

4. Kulikovsky A, Serebryakova M, Bantysh O, Metlitskaya A, Borukhov S, Severinov K, Dubiley S. 2014. The molecular mechanism of aminopropylation of peptide-nucleotide antibiotic microcin C. J Am Chem Soc 136:11168–11175.

5. Severinov K, Nair SK. 2012. Microcin C: biosynthesis and mechanisms of bacterial resistance. Future Microbiol 7:281–289.

6. Metlitskaya A, Kazakov T, Kommer A, Pavlova O, Praetorius-Ibba M, Ibba M, Krasheninnikov I, Kolb V, Khmel I, Severinov K. 2006. Aspartyl-tRNA synthetase is the target of peptide nucleotide antibiotic Microcin C. J Biol Chem 281:18033–18042.

7. Kazakov T, Vondenhoff GH, Datsenko KA, Novikova M, Metlitskaya A, Wanner BL, Severinov K. 2008. Escherichia coli peptidase A, B, or N can process translation inhibitor microcin C. J Bacteriol 190:2607–2610.

8. Novikova M, Kazakov T, Vondenhoff GH, Semenova E, Rozenski J, Metlytskaya A, Zukher I, Tikhonov A, Van Aerschot A, Severinov K. 2010. MccE provides resistance to protein synthesis inhibitor microcin C by acetylating the processed form of the antibiotic. J Biol Chem 285:12662–12669.

9. Tikhonov A, Kazakov T, Semenova E, Serebryakova M, Vondenhoff G, Van Aerschot A, Reader JS, Govorun VM, Severinov K. 2010. The mechanism of microcin C resistance provided by the MccF peptidase. J Biol Chem 285:37944–37952.

10. Krakowiak A, Pace HC, Blackburn GM, Adams M, Mekhalfia A, Kaczmarek R, Baraniak J, Stec WJ, Brenner C. 2004. Biochemical, Crystallographic, and Mutagenic Characterization of Hint, the AMP-Lysine Hydrolase, with Novel Substrates and Inhibitors. J Biol Chem 279:18711–18716.

11. Wang J, Fang P, Schimmel P, Guo M. 2012. Side chain independent recognition of aminoacyl adenylates by the Hint1 transcription suppressor. J Phys Chem B 116:6798–6805.

12. Brenner C. 2014. Histidine Triad (HIT) Superfamily, p. 1–8. In eLS. John Wiley & Sons, Ltd, Chichester, UK.

13. Brenner C. 2002. Hint, Fhit, and GalT: function, structure, evolution, and mechanism of three branches of the histidine triad superfamily of nucleotide hydrolases and transferases. Biochemistry 41:9003–9014.

14. Kijas AW, Harris JL, Harris JM, Lavin MF. 2006. Aprataxin forms a discrete branch in the HIT (Histidine Triad) superfamily of proteins with both DNA/RNA binding and nucleotide hydrolase activities. J Biol Chem 281:13939–13948.

15. Liu H, Rodgers ND, Jiao X. 2009. The scavenger mRNA decapping enzyme DcpS is amember of the HIT family of pyrophosphatases 21:4699–4708.

16. Holden HM, Rayment I, Thoden JB. 2003. Structure and Function of Enzymes of the Leloir Pathway for Galactose Metabolism. J Biol Chem 278:43885–43888.

17. Chou T-FF, Bieganowski P, Shilinski K, Cheng JJ, Brenner C, Wagner CR. 2005. 31P NMR and genetic analysis establish hinT as the only Escherchia coli purine nucleoside phosphoramidase and as essential for growth under high salt conditions. J Biol Chem 280:15356–15361.

18. Holland IB, Peherstorfer S, Kanonenberg K, Lenders M, Reimann S, Schmitt L. 2016. Type I Protein Secretion-Deceptively Simple yet with a Wide Range of Mechanistic Variability across the Family. EcoSal Plus 216; doi:10.1128/ecosalplus.ESP-0019-2015:0019-2015.

19. Bardaweel S, Pace J, Chou T-F, Cody V, Wagner CR. 2010. Probing the Impact of the echinT C-Terminal Domain on Structure and Catalysis. J Mol Biol 404:627–638.

20. Shah R, Maize KM, Zhou X, Finzel BC, Wagner CR. 2017. Caught before Released: Structural Mapping of the Reaction Trajectory for the Sofosbuvir Activating Enzyme, Human Histidine Triad Nucleotide Binding Protein 1 (hHint1). Biochemistry 56:3559–3570.

21. Liang G, Webster CE. 2017. Phosphoramidate hydrolysis catalyzed by human histidine triad nucleotide binding protein 1 (hHint1): A cluster-model DFT computational study. Org Biomol Chem 15:8661–8668.

22. Zhou X, Chou TF, Aubol BE, Park CJ, Wolfenden R, Adams J, Wagner CR. 2013. Kinetic mechanism of human histidine triad nucleotide binding protein 1. Biochemistry 52:3588–3600.

23. Waterhouse A, Bertoni M, Bienert S, Studer G, Tauriello G, Gumienny R, Heer FT, de Beer TAP, Rempfer C, Bordoli L, Lepore R, Schwede T. 2018. SWISS-MODEL: homology modelling of protein structures and complexes. Nucleic Acids Res 46:W296–W303.

24. Baugh L, Phan I, Begley DW, Clifton MC, Armour B, Dranow DM, Taylor BM, Muruthi MM, Abendroth J, Fairman JW, Fox D, Dieterich SH, Staker BL, Gardberg AS, Choi R, Hewitt SN, Napuli AJ, Myers J, Barrett LK, Zhang Y, Ferrell M, Mundt E, Thompkins K, Tran N, Lyons-Abbott S, Abramov A, Sekar A, Serbzhinskiy D, Lorimer D, Buchko GW, Stacy R, Stewart LJ, Edwards TE, Van Voorhis WC, Myler PJ. 2015. Increasing the structural coverage of tuberculosis drug targets. Tuberculosis (Edinb) 95:142–148.

25. Dolot R, Ozga M, Włodarczyk A, Krakowiak A, Nawrot B. 2012. A new crystal form of human histidine triad nucleotide-binding protein 1 (hHINT1) in complex with adenosine 5’-monophosphate at 1.38 Å resolution. Acta Crystallogr Sect F Struct Biol Cryst Commun 68:883–888.

26. Bantysh O, Serebryakova M, Zukher I, Kulikovsky A, Tsibulskaya D, Dubiley S, Severinov K. 2015. Enzymatic Synthesis and Functional Characterization of Bioactive Microcin C-Like Compounds with Altered Peptide Sequence and Length. J Bacteriol 197:3133–3141.

27. Chou T-F, Baraniak J, Kaczmarek R, Zhou X, Cheng J, Ghosh B, Wagner CR. 2007. Phosphoramidate pronucleotides: a comparison of the phosphoramidase substrate specificity of human and Escherichia coli histidine triad nucleotide binding proteins. Mol Pharm 4:208–217.

28. Chou T-F, Sham YY, Wagner CR. 2007. Impact of the C-terminal loop of histidine triad nucleotide binding protein1 (Hint1) on substrate specificity. Biochemistry 46:13074–13079.

29. Ho SN, Hunt HD, Horton RM, Pullen JK, Pease LR. 1989. Site-directed mutagenesis by overlap extension using the polymerase chain reaction. Gene 77:51–59.

30. Marchler-Bauer A, Bo Y, Han L, He J, Lanczycki CJ, Lu S, Chitsaz F, Derbyshire MK, Geer RC, Gonzales NR, Gwadz M, Hurwitz DI, Lu F, Marchler GH, Song JS, Thanki N, Wang Z, Yamashita RA, Zhang D, Zheng C, Geer LY, Bryant SH. 2017. CDD/SPARCLE: functional classification of proteins via subfamily domain architectures. Nucleic Acids Res 45:D200–D203.

31. Schaffer AA. 2001. Improving the accuracy of PSI-BLAST protein database searches with composition-based statistics and other refinements. Nucleic Acids Res 29:2994–3005.

32. Edgar RC. 2010. Search and clustering orders of magnitude faster than BLAST. Bioinformatics 26:2460–2461.

33. Edgar RC. 2004. MUSCLE: A multiple sequence alignment method with reduced time and space complexity. BMC Bioinformatics 5:1–19.

34. Söding J. 2005. Protein homology detection by HMM-HMM comparison. Bioinformatics 21:951–960.

35. Price MN, Dehal PS, Arkin AP. 2010. FastTree 2--approximately maximum-likelihood trees for large alignments. PLoS One 5:e9490.

36. Chou T-FF, Wagner CR. 2007. Lysyl-tRNA synthetase-generated lysyl-adenylate is a substrate for histidine triad nucleotide binding proteins. J Biol Chem 282:4719–4727.

37. Zukher I, Pavlov M, Tsibulskaya D, Kulikovsky A, Zyubko T, Bikmetov D, Serebryakova M, Nair SK, Ehrenberg M, Dubiley S, Severinov K. 2019. Reiterative Synthesis by the Ribosome and Recognition of the N-Terminal Formyl Group by Biosynthetic Machinery Contribute to Evolutionary Conservation of the Length of Antibiotic Microcin C Peptide Precursor. MBio 10: e00768–19.

